# Accelerating ligand discovery by combining Bayesian optimization with MMGBSA-based binding affinity calculations

**DOI:** 10.1101/2025.06.22.660936

**Authors:** Lucas Andersen, Max Rausch-Dupont, Alejandro Martínez León, Andrea Volkamer, Jochen S. Hub, Dietrich Klakow

## Abstract

Predicting protein–ligand binding affinity with high accuracy is critical in structure-based drug discovery. While docking methods offer computational efficiency, they often lack the precision required for reliable affinity ranking. In contrast, molecular dynamics (MD)-based approaches such as MMGBSA provide more accurate binding free energy estimates but are computationally intensive, limiting their scalability. To address this trade-off, we introduce an active learning framework that automates molecule selection for docking and MD simulations, replacing manual expert-driven decisions with a data-efficient, model-guided strategy. Our approach integrates fixed — partly pre-trained deep learning — molecular embeddings (MolFormer, ChemBERTa-2, and Morgan fingerprints) with adaptive regression models (e.g. Bayesian Ridge and Random Forest) to iteratively improve binding affinity predictions. We evaluate this approach retrospectively on a new dataset of 59,356 chemically diverse compounds from ZINC-22 targeting the MCL1 protein using both AutoDock Vina and MMGBSA binding free energy scores. Our results show that incorporating MMGBSA scores into the active learning loop significantly enhances performance, recovering 79.9% of the top 1% binders in the whole dataset, compared to only 6.7% when using docking scores alone. Notably, MMGBSA exhibits a stronger correlation with experimental binding affinities than AutoDock Vina on our dataset and enables more accurate ranking of candidate compounds in a runtime efficient way. Furthermore, we demonstrate that a one-at-a-time acquisition active learning strategy consistently outperforms traditional batched acquisition, the latter achieving just 78.4% recovery with MolFormer and Bayesian Ridge. These findings underscore the potential of integrating deep learning-based molecular representations with MD-level accuracy in an active learning framework, offering a scalable and efficient path to accelerate virtual screening and improve hit identification in drug discovery.

## 1 Introduction

Drug discovery is a lengthy and costly process, requiring approximately 12 years and an investment of approximately $1.8 billion USD.^1^ This process encompasses several challenges along the drug development pipeline, from target identification over hit compound finding, to hit-to-lead optimization and ultimately to (pre)clinical trials. Once a target is identified, promising (bio)active compounds are searched through experimental high-throughput screening (HTS) or its computational counterpart, virtual screening (VS). Although VS is generally less accurate than HTS, it allows a rapid evaluation of a much broader chemical space. ^2^

To facilitate the identification of promising compounds, several extensive screening libraries are available. The ZINC22^3^ database contains over 50 billion chemical compounds that can be ordered from commercial suppliers. Another example is Enamine’s Real database^4^ featuring building blocks that can be combined to 70 billion compounds, which can be synthesized on demand with a success rate of 80% within two weeks.

Different computational methods have been developed to search the vast chemical space, which vary in their computational cost and in their accuracy in correctly ranking ligands by binding affinity. Efficient molecular docking algorithms have been used with the aim to predict the optimal binding conformation of ligands within a protein binding pocket by minimizing an affinity scoring function^5,6^ and as enrichment tool to prioritize ligands that are more likely to bind to a target. However, docking protocols typically neglect the flexibility of the protein and rely on highly simplified models for protein-ligand interactions and solvent contributions, which may lead to poor correlations between docking scores and experimental binding affinities.^7^ In contrast, methods based on molecular dynamics (MD) simulations can account for protein flexibility and explicit solvent effects, but they come at a significantly higher computational cost compared to docking. Among MD-based methods, free energy perturbation (FEP) may be considered as a gold standard for affinity predictions as it relies on a rigorous statistical framework along an alchemical binding pathway and treats the solvent explicitly throughout the simulations and affinity calculations. Simplified MD-based approaches are given by end-point free energy techniques such as the Molecular Mechanics Generalized-Born Surface Area (MMGBSA) method. MMGBSA involves explicit-solvent simulations of both, the bound and the unbound state, however it approximates the solvent and entropy contributions to the binding affinity with implicit models.^8–10^ In this study, we score ligands using MMGBSA, an MD-based binding affinity estimation technique^11^, and AutoDock Vina^5^, a tool built upon an empirical scoring function.

Active learning approaches, such as Bayesian Optimization^12^, can build upon binding affinities from MD-based methods such as MMGBSA, enabling binding affinity predictions with MD-level accuracy while maintaining scalability for large-scale VS applications. In active learning, binding affinity predictions are performed iteratively, thereby avoiding the calculation of binding affinities for all available molecules, and instead allowing the algorithm to focus on promising regions of the chemical space. During each iteration, a surrogate model selects a subset of molecules for simulation. The molecules are selected such that the likelihood of finding the molecule with the lowest binding free energy in the entire database is maximized. After each iteration, the surrogate model is updated with the newly computed binding affinities obtained from the simulation, thereby refining its predictions for subsequent rounds.

While the optimization of lead candidates using generative models has gained significant attention in recent years^13–16^, active learning^12^ has proven highly effective for screening large datasets in drug design campaigns. ^17–23^ Graff *et al*. ^21^ analyzed several surrogate models in a pool-based active learning setting and found that the best performance is achieved when using multilayer perceptrons (MLP) and message-passing neural networks^24^ (MPNN) as surrogate models. However, training such surrogate models is time-consuming. Cao *et al*. ^25^ proposed a method in which the entire model, including the pretrained components, is updated during each training iteration. While continuing to train a pretrained model can enhance performance, it is computationally expensive and requires careful handling to prevent the new updates from overwriting previously learned knowledge.^26^

In this work, we demonstrate how combining Bayesian active learning with binding affinities from MD-based MMGBSA calculations can both accelerate the drug discovery process and bring MD-level accuracy to virtual screening applications. To this end, we compiled a large data set of binding affinities for 59,356 chemically diverse compounds from the ZINC-22 database, targeting the binding pocket of the myeloid cell leukemia 1 (MCL1) protein. MCL1 is a promising target for anti-cancer therapies that has been used previously to benchmark binding affinity calculations^27,28^.

We show that by using embeddings from a pretrained chemical language model without further finetuning, the surrogate model can be a simple model such as ordinary linear regression while maintaining the high level of performance. Additionally, retraining the model speeds up the active learning process during the early stage of learning. Specifically, we find that using a pretrained embedding model does not require retraining the full model, instead updating a simple linear regression model after each iteration yields the same retrieval of top binders and also offers the possibility of one-at-a-time molecule acquisition. Using our active learning pipeline, we obtain 79.9% of the top-1% binders according to MMGBSA after querying 6% of the approximately 60,000 compounds. In addition, we find that learning docking scores is easier than learning MMGBSA scores, as shown by retrieving 97.1% of the top-1% binders according to docking scores after querying 6% of the compounds. However, since MMGBSA scores exhibit a much stronger correlation with experimental binding affinities compared to docking scores, our results suggest that the combination of active learning with MMGBSA scoring provides a compelling balance between computational efficiency and predictive accuracy for identifying promising ligand candidates.

## 2 Computational Background

### 2.1 Binding affinity calculations with MMGBSA

MMGBSA is an end-point free energy method that estimates binding affinities based on MD simulations of only the bound and unbound states. MMGBSA employs a thermodynamic cycle to estimate the binding affinity Δ`G` (Fig. 1c), leading to the definition:

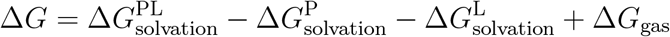

**Figure 1.**
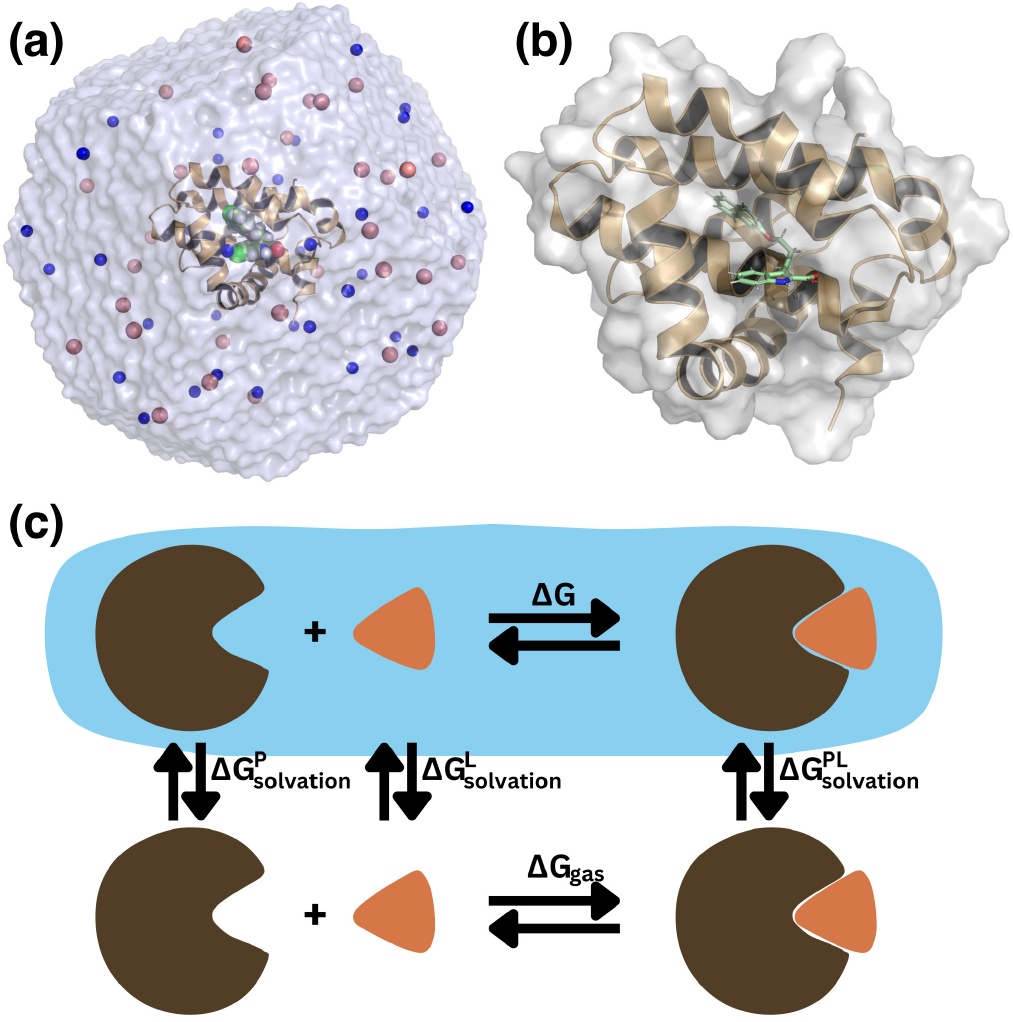
(a) MD simulations system of MCL1–ligand complex solvated in explicit solvent. The ligand’s smiles string is: `[O-]C(=O)c1[nH]c2ccccc2c1CCCOc1cc2ccccc2cc1`. The protein is shown as cartoon representation, the ligand as spheres, water as transparent surface, and NaCl ions as blue/red spheres. (b) Close-up view on the protein with bound ligand. (c) Thermodynamic cycle used for computing the binding free energy Δ*G for a ligand (orange) b*inding to a protein (brown) using MMGBSA. The blue area illustrates states solvated in water. For definition of the the mathematical symbols, see text.

Here, ΔG 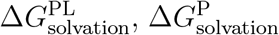 and 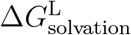 denote the solvation free energies of the protein-ligand complex, the protein, and the ligand, respectively, computed with the Generalized Born (GB) method or the Poisson-Boltzmann (PB) method^29^, together with the solvent-accessible surface area (SA). These methods operate within the framework of implicit solvent models, simplifying the representation of solvation effects.

Δ*G*_*gas*_ *denotes* the binding affinity in the gas phase, which is computed as the difference in expected values of potential energy, derived from a molecular mechanics force field, between the products (protein-ligand complex) and the reactants (protein and ligand). This term may optionally include an entropic correction contribution.^11,30^ While the MD simulations are typically conducted in explicit solvent (Fig. 1a) to ensure realistic conformational sampling, the solvation free energy terms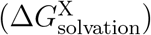) are estimated using implicit solvation models (GB or PB combined with SA). Although absolute binding free energy values from MMGBSA often deviate significantly from experimental binding affinities, the method has been shown to provide reasonably accurate ranking of ligands binding to the same target protein. ^27,31–33^

### 2.2 Bayesian Optimization

Bayesian Optimization (BO)^12^ is a technique for efficiently identifying the global optimum of a computationally expensive black-box function *f, w* here neither gradient information nor an analytical expression is available. Given a dataset of labeled inputs 𝒟, BO mitigates the cost of labeling the entire domain by constructing a probabilistic surrogate model 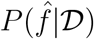 of the objective function *f*, which subsequently evaluates the utility of labeling each unlabeled data point. Since the objective is to identify the optimum of *f*, the utility is typically defined as a measure of improvement relative to the currently labeled data points. Gaussian Processes (GP)^34^ have traditionally been a popular choice for surrogate modeling, while Bayesian Neural Networks (BNN)^35,36^ have emerged as a more recent alternative.

Surrogate models are typically Bayesian for two main reasons. First, in theory, Bayes’ theorem allows efficient model updates when new labels are acquired, eliminating the need for costly full-model retraining. Second, Bayesian models provide access to the full posterior distribution, enabling more robust selection of the next evaluation point by considering, for example, the expected prediction rather than only the most likely prediction.

The utility of a new point can be assessed by acquisition functions such as *expected improvement* EI(*x)*.^37^ This criterion assigns zero utility to points that are not expected to improve upon the best observed value, *f* ^*⋆*^, while assigning the expected improvement to all other points. If the objective is to find a minimizer of *f*, EI is given by

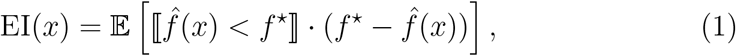

*W*here ⟦ · ⟧ denotes the Iverson bracket, which evaluates to 1 if its argument is true and otherwise to 0. The expectation is computed over all surrogate models 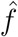 that have a non-zero posterior probability.

Assuming a Gaussian posterior distribution, the expected improvement permits the following closed-form expression:

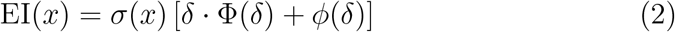

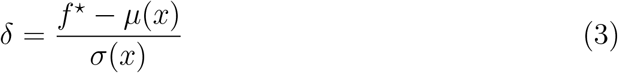

Here, Φ and *ϕ* denote the cumulative distribution and probability density function of the standard normal distribution, respectively, and *µ, σ* denote the mean and standard deviation of the posterior distribution at point *x.* Essentially, the first term assigns high values to points that are likely to improve (exploit), while the second term assigns higher values to points associated with a large uncertainty (explore). Thus, EI automatically balances exploitation and exploration.

#### 2.2.1 Bayesian Optimization with foundation models

In recent years, self-supervised pretraining of deep neural networks, such as transformer models^38^, has proven effective for learning rich input representations, particularly in fields like natural language processing ^39,40^, computer vision^41–44^, and cheminformatics^45–49^. Additionally, these pretrained models are highly effective for transfer learning, as their parameters already capture general data features, allowing them to adapt to downstream tasks with few labeled examples.^49–52^

The main difficulty in adapting such models to BO is that they are not Bayesian by default. Simple approaches, such as Monte Carlo Dropout^53^, cannot be easily applied to pretrained models that were not originally trained with dropout, as these models lack the necessary dropout layers to enable uncertainty estimation. To overcome this issue, we perform Bayesian linear regression, with the pretrained embeddings as input. Bayesian linear regression allows direct access to the posterior distribution, including uncertainty estimates. This approach has been shown to be a viable alternative to fully probabilistic methods. ^36,54^

## 3 Dataset

To simulate a realistic early-stage drug discovery scenario and evaluate our active learning approach, we construct a dataset as a representative subset of a large molecular library, such as ZINC.^3^ Starting from a few known binders, we have assembled a screening set by applying a loose similarity filter (MCS similarity with a threshold of 0.4) to ensure some chemical relevance while maintaining diversity. It is to note that despite the similarity-based selection criterion, the dataset of approximately 60,000 molecules is highly divers with many compounds differing substantially in their scaffold and functional groups.

### Protein target

As a case study, we select the MCL1 protein, which has been shown being a promising target for anti-cancer therapy.^55^ The crystal structure was taken from the protein data bank (PDB code 4HW3).^56^ Friberg *et al*. ^56^ provide experimental measurements of the inhibitory constant for 41 MCL1 binders.

### Small molecule data set

The ZINC22 compound collection, comprising more than 54 billion molecules, serves as screening library in our study.^3^ We compare each of these 54 billion molecules with the strongest experimental binder according to Friberg *et al*. ^56^ using a Tanimoto similarity coefficient. We measure similarity using the maximum common substructure (MCS) metric.^57^ We leverage RDKit^58^ to calculate the MCS and include all molecules above a similarity threshold of 0.4. Despite screening for similarity, most compounds show a low similarity to the query and the average pairwise similarity in the dataset is just 0.22. A histogram of the similarity distribution can be found in the supplementary material (Fig. S1). Moreover, the structural diversity of the dataset is further supported by the observation that many compounds achieve a significantly worse MMGBSA score than the experimentally validated binders (Fig. 2c).

**Figure 2.**
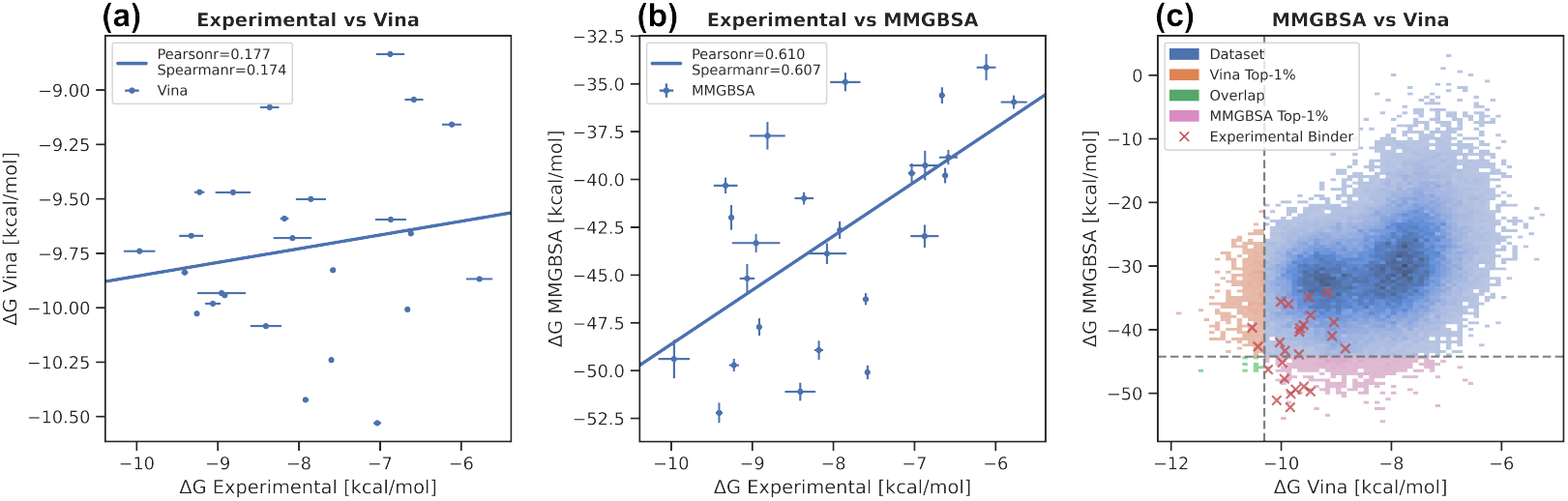
Comparison of binding affinities from MMGBSA or AutoDock Vina with experimental data. (a) Correlation between experimental binding affinity and AutoDock Vina scores for the set of experimentally known binders. Each ligand was docked inside of the protein with the search box spanning the MCL1 binding pocket. The RMSD values computed from the maximum common substructure between each ligand and the reference ligand of the crystal structure are provided in the Supplementary Data. (b) Correlation between experimental and MMGBSA-based binding affinity for the set of experimentally known binders. The docked poses as identified for (a) were used as starting structures for the MD simulations associated with the MMGBSA computation. (c) Correlation between AutoDock Vina and MMGBSA for the augmented dataset highlighting the top 1% of molecules according to the respective computational method. The red crosses indicate the computed score for the experimental binders. Each ligand was docked following the same procedure as in (a).

Afterwards, to obtain an initial pose for the MMGBSA computation we dock all molecules using AutoDock Vina. ^5^ For each ligand, we carry out MD simulations to obtain the binding free energy using MMGBSA (see Methods). The final size of our dataset comprises 59, 356.

Figures 2a/b show the correlation of the experimental binding affinities with MMGBSA and AutoDock Vina for the subset of experimentally known binders. The Vina score exhibits poor Pearson correlation with the experimental binding affinities (0.177) and poorly ranks the ligands as indicated by a Spearman correlation coefficient of 0.174 (Fig. 2a). As expected, MMGBSA overestimates the absolute binding affinity; nevertheless, it demonstrates substantially improved agreement with the experimental data, with Pearson and Spearman correlations of 0.610 and 0.607, respectively (Fig. 2b). Using MMPBSA instead of MMGBSA further improves the correlation with the experimental data with Pearson and Spearman correlations of 0.689 and 0.684, respectively (Fig. S7). These findings support our assumption that the computationally more demanding MMGBSA method provides better ligand ranking than AutoDock Vina. Notably, among the 59,356 compounds of our augmented data set, docking and MMGBSA scores correlate poorly (Fig. 2c), demonstrating that the MMGBSA and Vina scores are sensitive to different molecular properties. The orange and pink areas of Fig. 2c highlight the top 1% of molecules based on the MMGBSA score and the Vina score, respectively, demonstrating that the set of top-1% binders according to MMGBSA and Vina are nearly non-overlapping. Only 1.6% of the top-1% binders according to MMGBSA also belong to the top-1% binders according to the Vina score. Overall, MMGBSA correlates more strongly with the experimental binding data subset and identifies a largely distinct set of top binders from Vina. These results suggest that Vina may not reliably distinguish the true highest-affinity compounds, making MMGBSA the preferred choice despite its higher computational cost.

### 3.1 Datasets in the literature

To compare our active learning pipeline with previous work, we also benchmark against the dataset utilized by Graff *et al*. ^21^ and Cao *et al*. ^25^ . The dataset comprises docking scores for the 50,240 molecules taken from the Enamine’s Discovery Diversity Collection (`Enamine50k`), docked against thymidylate kinase using AutoDock Vina, as described by Graff *et al*. ^21^ . Unlike previous studies, our work additionally incorporates MMGBSA scores.

## 4 Methods

### 4.1 Binding free energy calculations using MMGBSA

MMGBSA and MMPBSA calculations were prepared, executed, and analyzed using our recently published BindFlow pipeline^27^. Details of the simulations are described in the following sections.

#### Protein-Ligand complex generation

The crystal structure of MCL1 was taken from the Protein Data Bank (PDB: 4HW3^56^). Starting with the SMILES representation of the ligands, RDKit^58^ and EasyDock^59^ were used to obtain PDBQT files of the ligands. Ligands were protonated using Dimorphite-DL^60^ at pH 7.4. Ligands were docked into the MCL1 binding pocket with AutoDock Vina^5^, using a cubic box centered at the reference ligand location at (67.9, − 32.6, 27.2) Å and using a box length of 20 Å. In case that Dimorphite-DL proposed more than one protonation state, the state with strongest affinity according to the Vina score was used for follow-up simulations. An example conformation of a ligand in the MCL1 pocket is shown in Fig. 1a/b.

#### MD simulation

The protein-ligand complex was placed in an octahedral simulation box using a distance of at last 1.5 nm between the protein and the box surface. The system was solvated with water and neutralized with 150 mM of NaCl. Protein interactions were described by the Amber99sb-ildn force field^61^ and the TIP3P water model^62^ was applied. Ligand interactions were described with the Open Force Field, version 2.0.0.^63^ Ions were described with the Amber parameters. ^61^ The ligand topology file was generated with TOFF.^64^ Hydrogen mass repartitioning was used with a hydrogen mass factor of 3. Genheden and Ryde ^29^ recommend an equilibration time of 100 to 200 ps for MMGBSA calculations. In this study, we adopted a multi-step approach with a total equilibration time of 107.5 ps. Following energy minimization, the system was equilibrated through the following steps: (i) NVT-equilibration with an integration time step of Δ*t* = 2 fs for 10 ps; (ii) NVT-equilibration with Δ*t* = 3 fs for 15 ps; (iii) NPT-equilibration with Δ*t* = 3 fs for 22.5 ps; (iv) NPT-equilibration with Δ*t* = 4 fs for 60 ps. Position restraints on the heavy atoms using a force constant of 2500 kJ mol^−1^nm^−2^ were activated from steps (i) to (iii), and removed for subsequent steps. Dispersive interactions and short-range repulsion were describe with a LennardJones potential with a cutoff at 1 nm. Electrostatic interactions were computed with the particle-mesh Ewald (PME) method using a real-space cutoff at 1 nm.^65,66^ Bond lengths involving hydrogen atoms were constrained with LINCS.^67^ The pressure was controlled using the Parrinello-Rahman barostat with a time constant of 2.0 ps.^68^ The temperature was controlled at 298.15 K using the Langevin thermostat with an inverse friction constant of 2.0 ps. During production simulations, a time step of 4 fs was used.

#### MMGBSA calculation

After equilibration, the system was simulated for 950ps. Frames were extracted every 50 ps, yielding 20 starting conformations for 20 production runs of 100 ps each. Both the 950 ps simulation and the 100 ps production runs were conducted under the same conditions as step (iv). MMGBSA-based binding affinities were computed from 20 frames extracted from each production run using gmx_MMPBSA.^11,69^ The final binding affinity is reported as the average and standard error across these 20 simulations. Figure 3 illustrates the MMGBSA computation pipeline. The GB-OBC2 model^70^ was used with an internal dielectric constant of 1.0 and external dielectric constant of 78.5, the mbondi2 radii set, a surface tension of 0.0072 k/mol/Å^2^, and a salt concentration of 150 mM. Following recommendations by Su *et al*. ^71^, the entropy term was not included. An example MD simulation system including topology and parameter files is provided on GitHub at https://github.com/uds-lsv/bayesian-optimization-mmgbsa.

**Figure 3.**
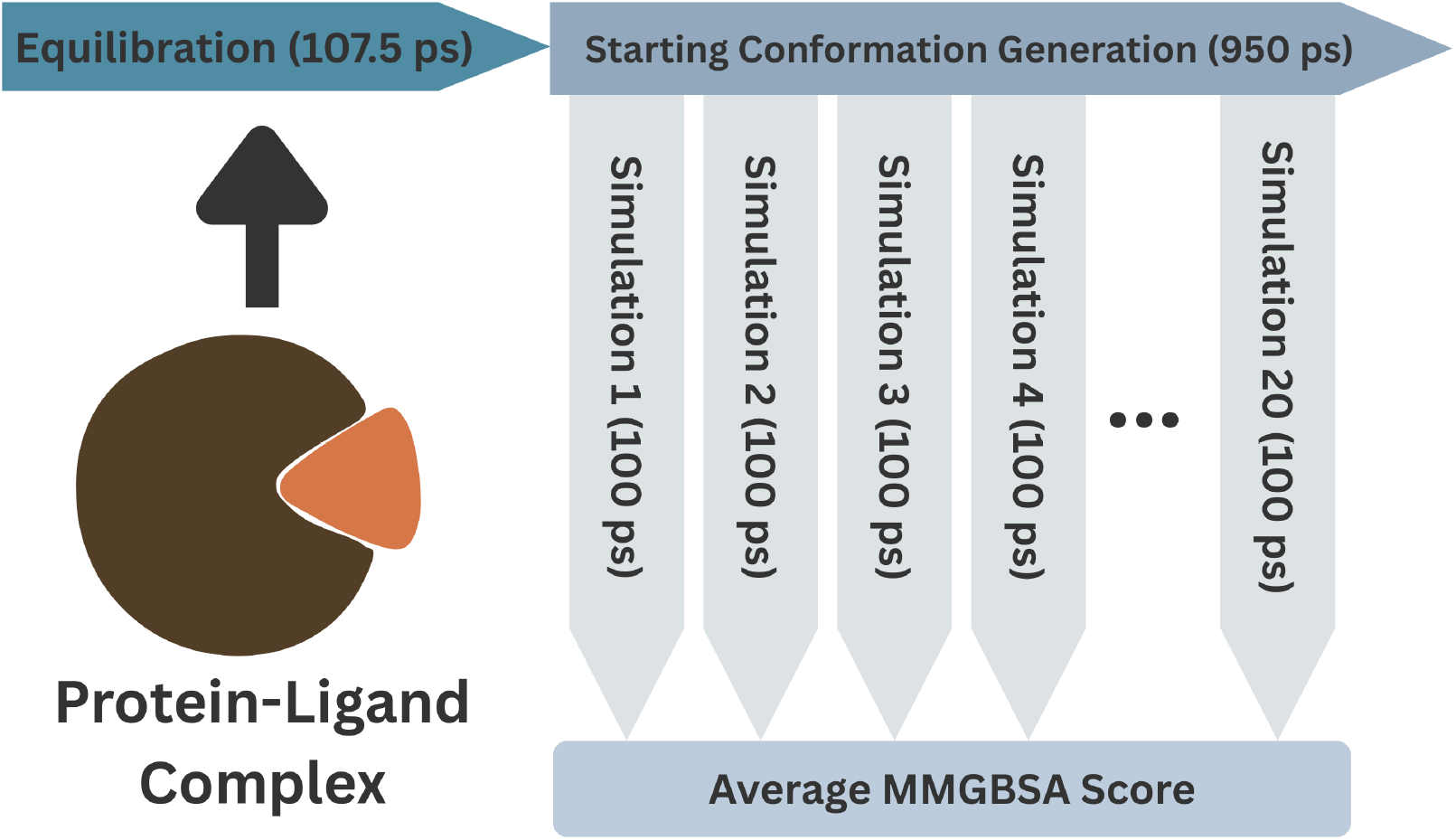
Pipeline of the MD simulation workflow: Each protein-ligand complex was equilibrated for 107.5 ps (25 ps NVT, 82.5 ps NPT). Then, 20 starting conformations were generated by extracting every 50 ps the conformation during a continuous MD simulation. Afterwards, each extracted conformation was used as the starting conformation for a 100 ps 100 ps production run, during which 20 frames were extracted. A total of 20 20 = 400 frames are generated. The final score is computed as the average MMGBSA score of each of the 20 ·100 ps runs. The total MD simulation time per protein-ligandcomplex amounts to 107.5 ps + 950 ps + 20 ·100 ps = 3057.5 ps ≈ 3 ns.

### 4.2 Active Learning pipeline

The pipeline is illustrated in Figure 4. The pipeline features the interplay of three central components: the surrogate model, the acquisition function, and the binding affinity computation technique. The surrogate model is the base AI model that is used, on the one hand, to predict binding affinities and, on the other hand, to estimate the uncertainty of the model prediction (Step (1) in Fig. 4). The surrogate model comprises two parts. The first part is an embedding method that transforms the molecules represented as a SMILES string to a high-dimensional vector, whereas the second part is a regression model that predict the binding affinity from the high-dimensional vector. We explored three different techniques to map the SMILES representation of a molecule to a high-dimensional vector: Morgan fingerprints ^72^, ChemBERTa2^47^, and MolFormer^48^. The latter two methods are transformer-based language models that have been pretrained using self-supervised learning.

**Figure 4.**
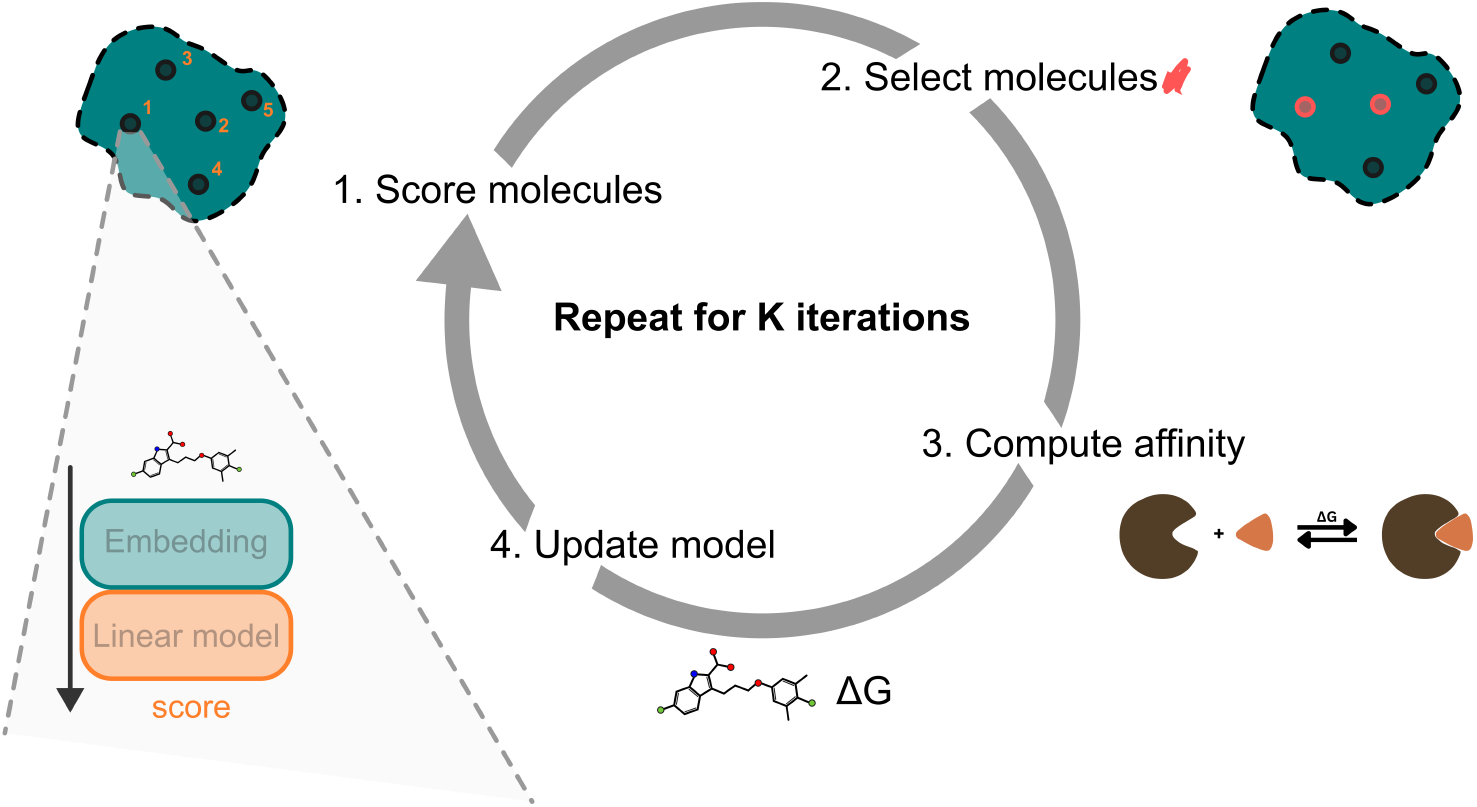
Pipeline of the active learning process: (1) All molecules are initially scored using the current surrogate model. (2) A subset of molecules is then selected with the acquisition function for binding affinity evaluation. (3) Binding affinities are computed using either classical docking methods or MD-based techniques such as MMGBSA. (4) The surrogate model is retrained with the additional data to refine predictions.

The acquisition function is a mathematical function that computes the informational value of a molecule based on the model prediction and uncertainty estimation. Step (2) of Fig. 4 uses the acquisition function to compute the informational value of each molecule and selects a predefined number of most valuable molecules for which the binding affinity is computed. After a selection of promising molecules, Step (3) computes the respective binding affinities, for instance using an MD-based technique such as MMGBSA, thereby expanding our dataset. Step (4) of Fig. 4 uses the expanded dataset to retrain our surrogate model for the next iteration. The process is repeated starting at Step (1).

In summary, in each iteration, we score all unlabeled molecules using the surrogate model and use the acquisition function to select the set of molecules with the highest utility value. In case we acquire not only one but a batch of *b* data points per iteration, we update the model only every *b* iterations. After the label is queried, the regression head of the surrogate model is retrained. The initial set of computed binding affinities comprises a single molecule taken as the molecule closest to the centroid in the embedding space. The pseudo-code of the active learning pipeline is shown in algorithm 1.

### 4.3 Bayesian linear regression

Bayesian Optimization requires access to the predictive posterior distribution of the surrogate model. We choose Bayesian linear regression as a simple model that fulfills this requirement. In combination with a pretrained embedding model, this is essentially equivalent to a Gaussian Process (GP) with a fixed basis function (the embedding model) and a dot product kernel.^34^ Compared to usual linear regression, Bayesian linear regression assumes a prior distribution over the weights. In particular, let ***w*** ∈ ℝ^*d*^ be the weights of the model, we assume *p(****w***|*α) = 𝒩* (0, *α*^*−1*^***I***), with *α*^*−1*^ being the variance. As is standard in regression tasks, we assume that the target is corrupted by independent Gaussian noise, i.e. *y(****x****) = f(****x***) + ϵ, where *p(ϵ) = 𝒩* (0, *β*^*−1*^*), f(****x****) =* ***w***^*T*^ ***x***, and **x** is the embedding vector of the ligand. Let ***X*** ∈ ℝ^*n×d*^ be the training inputs with associated targets **y** ∈ ℝ^*n*^. By Bayes theorem, the posterior distribution of the weights is a Gaussian with mean 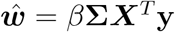and inverse covariance matrix **Σ**^−1^ = *α****I*** *+ β****X***^*T*^ ***X***. Marginalizing the posterior distribution over the weights yields the predictive posterior distribution as a Gaussian with mean 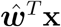and variance *σ*^*2*^(***x****) = β +* ***x***^*T*^ **Σx**. For BO applications, the true values of *α* and *β* are typically not known. We infer them by maximizing the marginal likelihood. ^73^ For more information we refer to Bishop ^73^ or Murphy ^74^ . Notably, the combination with a pretrained model is essentially the approach taken by Snoek *et al*. ^36^, and the weights of the embedding model may be considered as hyperparameters of the basis function.

#### Algorithm 1

Bayesian Optimization with Surrogate Model

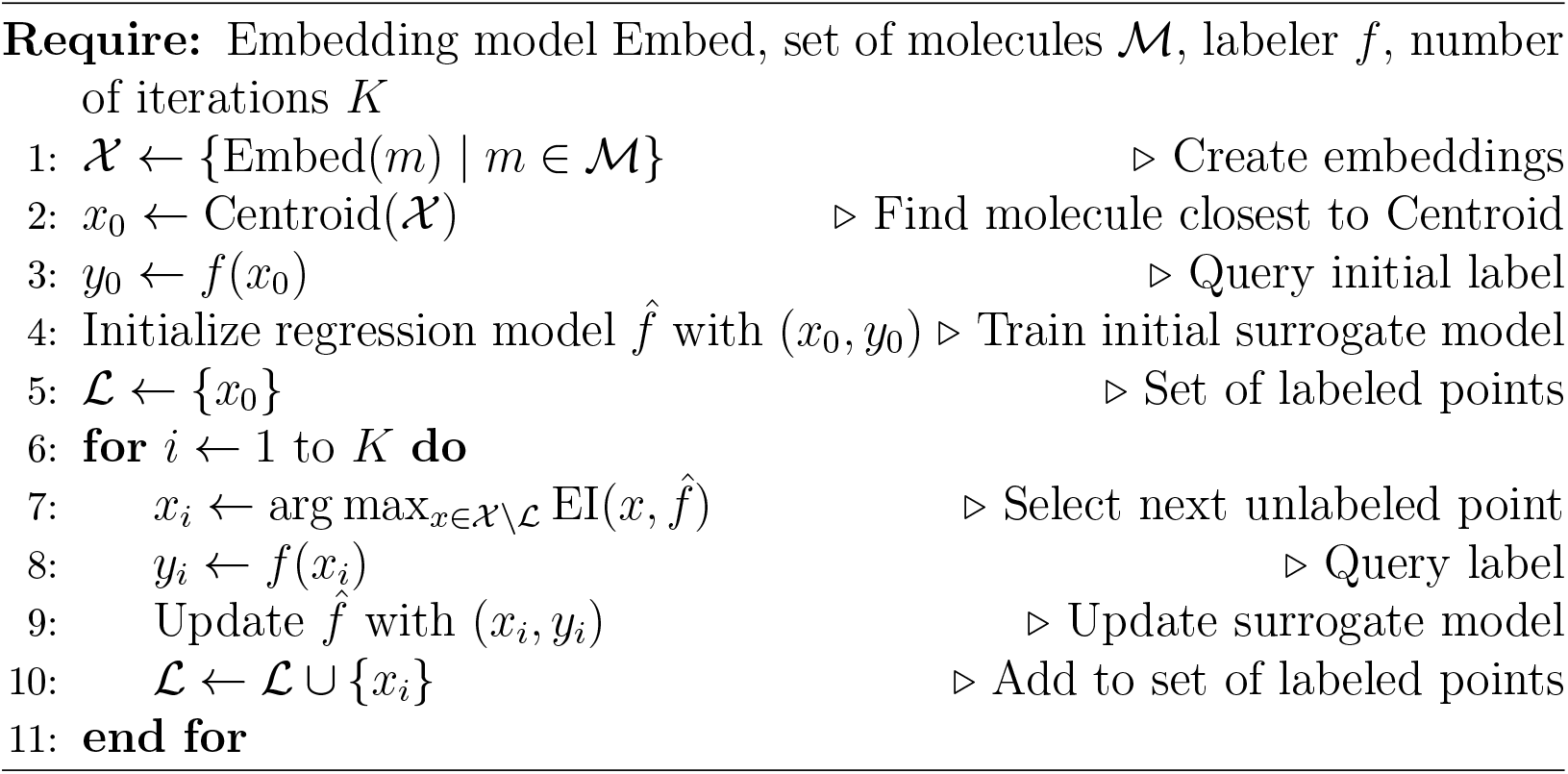

## 5 Results

As discussed in the dataset section, in the subset of experimentally tested binders, MMGBSA scores correlate more strongly with experimental binding affinities compared to Vina scores, suggesting that MMGBSA provides a more reliable binding affinity estimate within this subset (Fig. 2a/b). These differences are likely due to the explicit consideration of protein and ligand flexibility in the MD simulation. Notably, the sets of top-1% binders identified by MMGBSA and AutoDock Vina differ substantially in the full dataset. These results indicate that using docking scores alone may not reliably identify the highest-affinity compounds, whereas MMGBSA may provide a more consistent selection.

### 5.1 Updating only the regression model is sufficient for rapid retrieval of top binders

Prior work by Graff *et al*. ^21^ and Cao *et al*. ^25^ benchmarked the performance of a surrogate model in finding top-1% ligands among the `Enamine50k`. As scores, these authors used docking scores against thymidylate kinase obtained by Autodock Vina (see Datasets). To compare our pipeline with previous work, we follow the experimental setting by querying a batch of *b* molecules, where *b* is approximately 1% of the dataset size, corresponding to *b* = 500 for the `Enamine50k` dataset. We perform six rounds, such that only 6% of the whole dataset is screened. The results are summarized in Table 1 (first column).

**Table 1.** Top-1% retrieval rate on the `Enamine50k` dataset based on Autodock Vina docking scores against thymidylate kinase. The first column shows the retrieval rate upon querying a batch of 500 molecules per iteration for six iterations, totaling 3000 molecules. The second column presents the retrieval rate upon querying 3000 molecules individually, one after another. We report results for both the greedy and upper confidence bound (UCB) acquisition strategies from Cao *et al*. ^25^ (rows 1-4) and compare to the results obtained by our method when using expected improvement (rows 5-10)

|  | Batched Query | Single Molecule Query |
| --- | --- | --- |
| MolFormer (Greedy)<sup>25</sup> | 78.36 | **X** |
| MolFormer (UCB)<sup>25</sup> | 79.24 | **X** |
| RF (Greedy)<sup>25</sup> | 54.52 | **X** |
| RF (UCB)<sup>25</sup> | 37.08 | **X** |
| ChemBERTa-2 + Linear | 78.23 | 81.85 |
| ChemBERTa-2 + RF | 65.73 | 73.99 |
| MolFormer + Linear | 74.39 | 78.02 |
| MolFormer + RF | 40.52 | 58.26 |
| Morgan fingerprints + Linear | 65.12 | 73.79 |
| Morgan fingerprints + RF | 39.91 | 52.62 |

To select new molecules for evaluation, we score the molecule’s utility using the Expected Improvement (EI) criterion.^37^ Prior work has also considered alternative acquisition functions such as the Upper Confidence Bound (UCB) and surrogate model predictions (greedy selection).^21,25^ In contrast to UCB, EI does not require tuning of additional hyperparameters. Following Cao *et al*. ^25^, we assess the model performance using the retrieval rate of the top-1% of molecules according to the binding affinity score. Cao *et al*. ^25^ report a top-1% retrieval rate of 79.24% on `Enamine50k` using the MolFormer model and updating all parameters after querying each batch.^48^ In comparison, our approach combining MolFormer with a linear model (`MolFormer+Linear`) achieves a slightly lower rate of 74.39%. However, substituting MolFormer with ChemBERTa-2 embeddings (`ChemBERTa-2+Linear`) improves performance, yielding a retrieval rate of 78.23%. The similar performance of the batched acquisition and one-at-a-time acquisition models indicates that updating the regression model instead of updating the entire pipeline including the regression model and the much larger embedding model has minimal impact on the overall retrieval rate. Thus, it is not required to further train the embedding model. Furthermore, we find that using a Random Forest model instead of the linear regression model significantly worsens performance in all cases (Table 1). A similar trend is observed with Morgan fingerprints, which exhibit comparatively lower retrieval performance than models based on pretrained embeddings. Consequently, we focus our further analysis on results obtained using the ChemBERTa-2 and MolFormer embedding models in combination with the linear regression model. Additional results involving the RF model and Morgan fingerprints are presented in the supplementary material (S2–S4). In conclusion, these findings suggest that full model finetuning is not necessary; instead, updating only the regression model after each iteration is sufficient to achieve strong retrieval performance.

### 5.2 Querying one-at-a-time molecule instead of querying batches of molecules improves top-1% retrieval rate

Previous work has employed deep neural architectures, for which finetuning until convergence is computationally expensive.^21,25^ As a result, prior approaches queried large batches, typically around 1% of the dataset, in each iteration to reduce the number of model updates. By updating only the regression head of our surrogate models, training becomes significantly faster, making it feasible to use batch sizes as small as a single molecule per iteration. By querying a single molecule per iteration, we achieve a top-1% retrieval rate of 81.85% using `ChemBERTa-2+Linear` on the `Enamine50k` dataset, surpassing the previous best performance of 79.24% (Table 1, compare first with second column). The advantage of querying single molecules is more evident during the early stages of the active learning process (Fig. 5): here, top-1% retrieval rate increases more rapidly with a one-at-a-time acquisition as compared to querying batches of 500 molecules, irrespective of the choices for the surrogate and embedding model.

**Figure 5.**
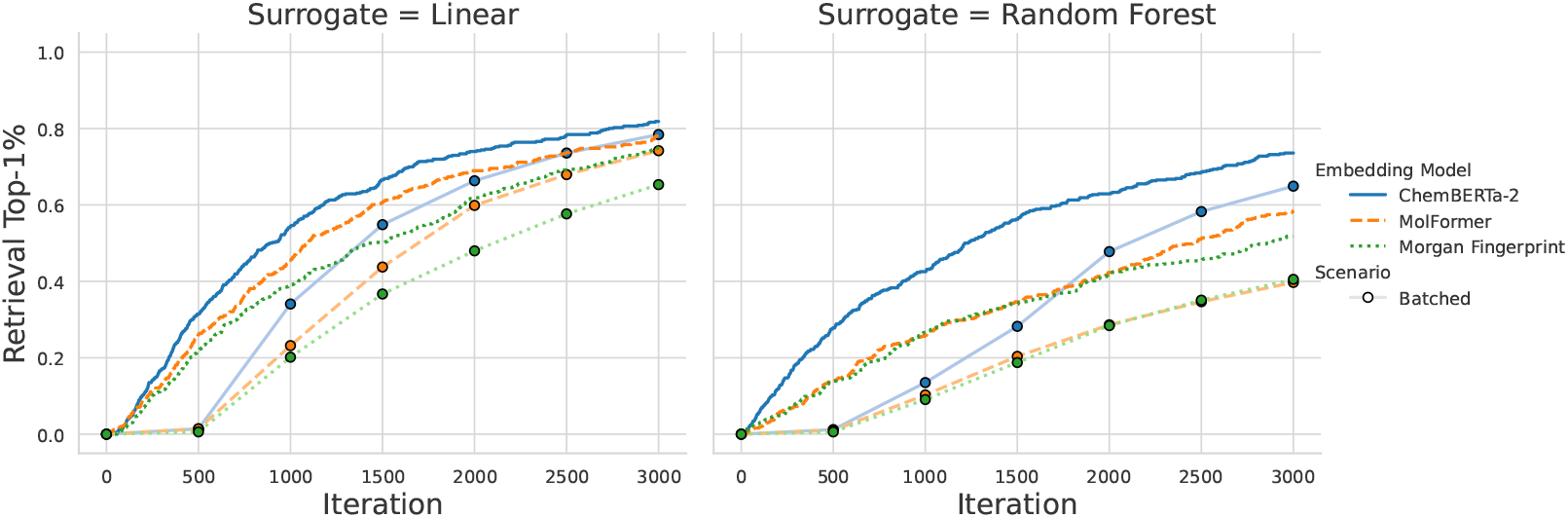
Top-1% retrieval rate based on docking scores of the `Enamine50k` dataset as a function of the number of screened molecules. Lines without markers represent one-at-a-time molecule acquisition, while lines with markers correspond to batch acquisitions of 500 molecules per iteration. Color codes and line styles indicate choice of the embedding model (see legend). Left: based a linear regression model. Right: Based on a Random Forest model.

### 5.3 Active learning with MMGBSA scores is more challenging than active learning with docking scores

Having established our active learning pipeline and compared it to previous work on the `Enamine50k` dataset, we next apply our pipeline to our newly derived dataset comprising 59,356 compounds taken from the ZINC22 database binding to MCL1 (see Datasets). Accordingly, we analyze the performance of active learning on both the docking scores (`MCL1-Docking`) and the MD-based MMGBSA binding affinity (`MCL1-MMGBSA`), while again comparing batched-query active learning approach with the one-at-a-time variant. Focusing first at the `MCL1-Docking` dataset, the batched model results in comparable scores between the MolFormer and ChemBERTa-2 models, both achieving over 90% retrieval rate (Table 2). Querying a single molecule, instead of a batch, greatly improves the top-1% retrieval during the early stages of the active learning process (Fig. 6, left) and, after 3600 iterations, improves the top-1% retrieval rate in both cases, with `MolFormer+Linear` reaching 95.8% and `ChemBERTa-2+Linear` reaching 97.1%. These advantages of the one-at-a-time acquisition are in line with the findings on the docking-based `Enamine50k` dataset presented in the last section. Furthermore, the finding that using MolFormer and ChemBERTa-2 yields similar learning rates aligns with the findings on the `Enamine50k` dataset.

**Table 2.** Top-1% retrieval rate of the linear regression model based on docking and MMGBSA scores for the MCL1 dataset. Full results, including retrieval rates using Morgan fingerprints and a random forest regression model, are provided in the supplementary material

| (a) MCL1-Docking |  |  |
| --- | --- | --- |
|  | Batched Query | Single Molecule Query |
| ChemBERTa-2 | 94.6 | 97.1 |
| MolFormer | 93.8 | 95.8 |

| (b) MCL1-MMGBSA |  |  |
| --- | --- | --- |
|  | Batched Query | Single Molecule Query |
| ChemBERTa-2 | 59.9 | 62.7 |
| MolFormer | 78.4 | 79.9 |

**Figure 6.**
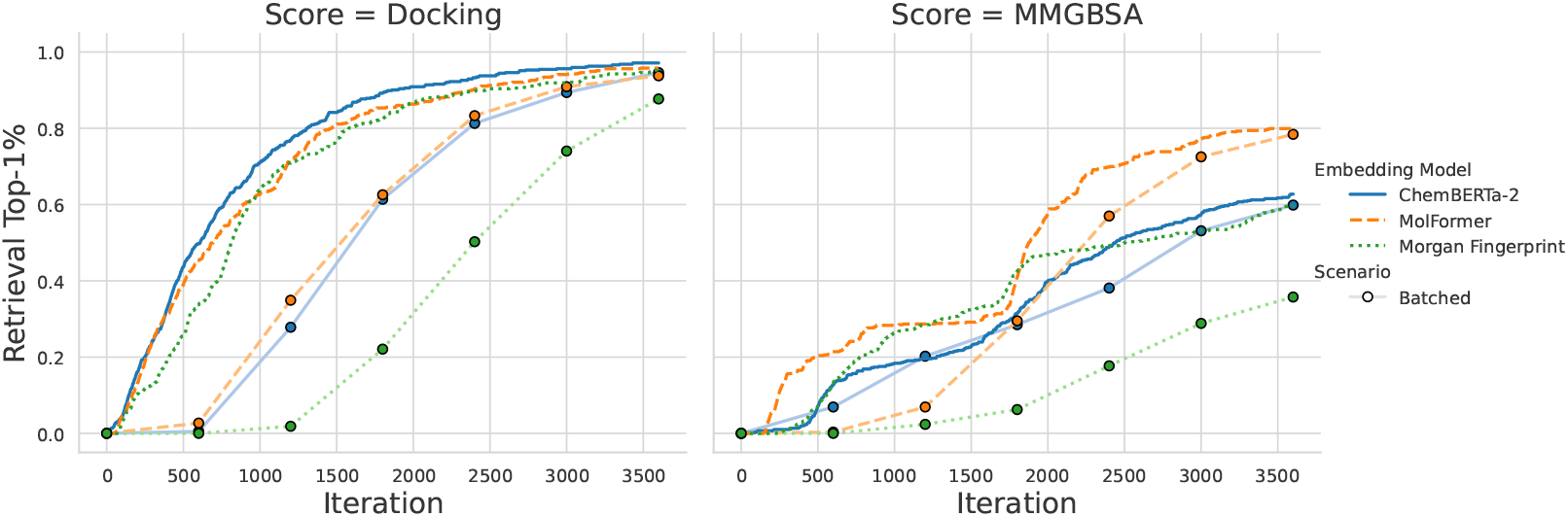
Top-1% retrieval rate of molecules based on docking and MMGBSA scores using a linear regression model. Lines without markers represent one-at-a-time molecule acquisition, while lines with markers correspond to batch acquisitions of 500 molecules per iteration. Results for the retrieval process using the random forest (RF) model are provided in the supplementary material .

Applying our pipeline to `MCL1-MMGBSA` dataset shows successful learning of MMGBSA scores. A key finding of our study is that the pipeline achieves top-1% retrieval rates of up to 79.9% upon querying 6% of the compounds when using single molecule query (SMQ) and MolFormer.

However, active learning of the `MCL1-MMGBSA` scores also revealed striking differences to the `MCL1-Docking` scores. We obtained lower top-1% retrieval rates when using MMGBSA scores (62.7% and 79.9%, SMQ) as compared to using docking scores (97.1% and 95.8%, SMQ; Table 2; Fig. 6, compare left with right panel). In addition, while the top-1% retrieval rate based on docking scores increases rapidly during early stages of the process, top-1% retrieval rate based on MMGBSA increases irregularly and requires far more queries before reaching significant retrieval rates. These findings suggest that learning docking scores is easier as compared to learning MMGBSA scores. We attribute these findings to the relative simplicity of the docking scoring algorithm, which we hypothesize renders the energy landscape smoother and its patterns easier for the model to learn. In contrast, MMGBSA scores may have a more complex structure owing to the physically more detailed representation of the ligand–protein interactions, rationalizing the need for a larger number of queries to achieve good top-1% retrieval rates.

As a first analysis in this direction, we compare by how much the activity score, either Vina or MMGBSA, between two points in embedding space changes. To this end, we compute pairwise differences, as well as the Euclidean distance between the datapoints in the MolFormer embedding space. Since the values of MMGBSA and Vina scores itself differ significantly in magnitude, we normalize the differences by their respective standard devia-tion. In Fig. 7 we show the average activity difference for a range of distance values. Even after normalization, we can observe that for essentially all distances, the MMGBSA scores show larger differences than the Vina scores. Indicating that, in the embedding space, the MMGBSA energy landscape is indeed less smooth than the Vina landscape. While our present analysis provides sufficient insight into the optimization process and learnability, a more detailed study of the effect of smoothness could offer valuable directions for future work.

**Figure 7.**
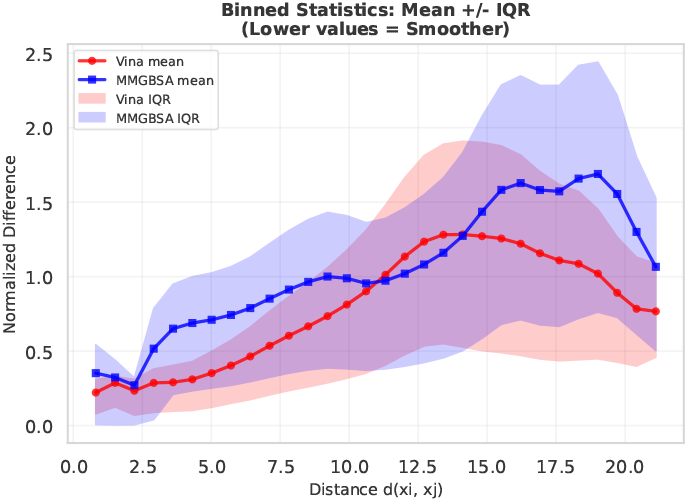
Average normalized activity score differences, binned by distance in MolFormer embedding space. Shaded areas represent the interquartile range (IQR). For most distances, the Vina score tends to show a smaller difference in the activity score, indicating less variation compared to the MMGBSA score.

Lastly, the choice of the embedding plays a different role during learning of MMGBSA scores as compared to docking scores (Fig. 6, right, blue and orange curves). Upon learning docking scores with the linear surrogate model, learning based on ChemBERTa-2 slightly outperforms learning based on MolFormer. In contrast, upon learning MMGBSA scores, using MolFormer leads to 20% better learning rates compared to using ChemBERTa2. To test whether these findings are consistent across independent active learning campaigns, we repeated training of our surrogate model, yet starting from different initial molecules (Fig. 8). Evidently, retrieval rates differ considerably upon using different initial molecules. However, top-1% retrievel rates after 3600 iterations using MolFormer consistently outperform the learning rates using ChemBERTa-2 or Morgan fingerprints, as shown by top-1% retrieval rates averaged over seven campaigns of 84%, 62%, and 50%, respectively. We hypothesize that the structural context embedded in MolFormer by the use of masked language modeling as a pretraining objective helps our surrogate model in learning MMGBSA scores. Thus, systematic testing of different embedding strategies for learning MMGBSA scores may be insightful in future studies.

**Figure 8.**
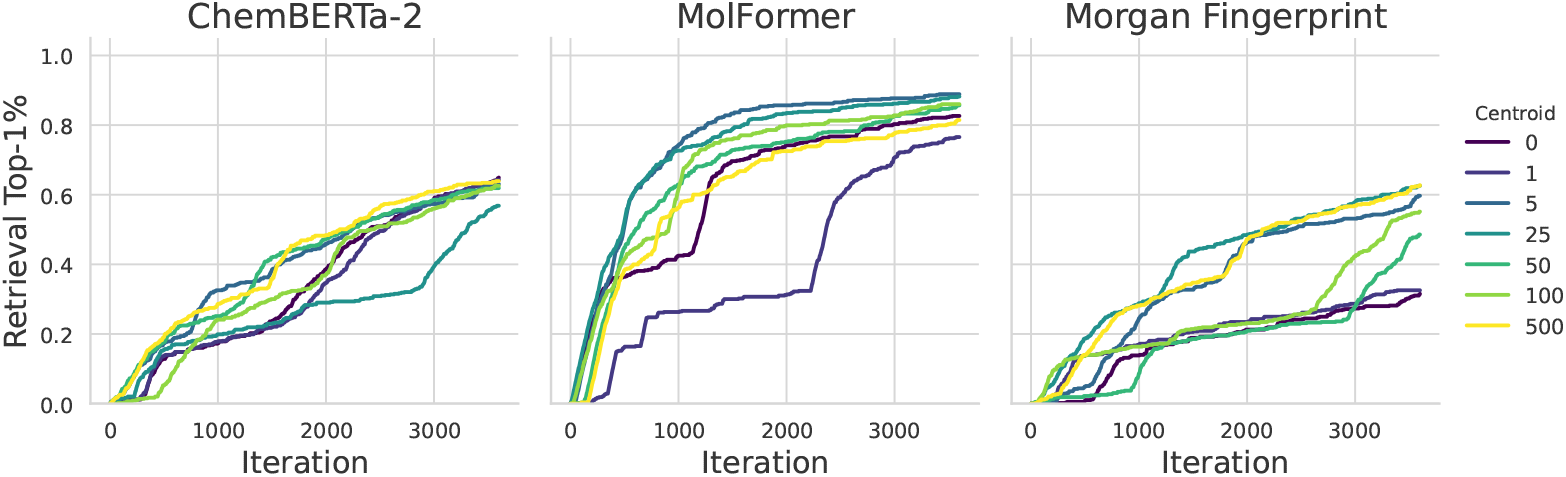
Top-1% retrieval rate of molecules based on MMGBSA scores using a linear regression, one-at-a-time acquisition, and different embedding models: from left to right, ChemBERTa-2, MolFormer, and Morgan fingerprints. Training of the surrogate model was initiated with different molecules highlighted by color taken as the centroid molecule (0), as well as the 1^st^, 5^th^, 25^th^, 50^th^, 100^th^ and 500^th^ nearest molecule to the centroid. Retrieval rates may differ considerably when using different initial molecules. However, Molfomer consistently outperforms both ChemBERTa-2 and Morgan fingerprints.

Considering that MMGBSA scores correlate more strongly with experimental binding affinities than docking scores and that its top binders differ significantly from those identified by docking scores (Fig. 2), we infer that MMGBSA-based selection may better identify candidates with stronger experimental binding affinities. While models may achieve higher top-1% retrieval rates when using docking scores, the binders selected according to the MMGBSA scores are likely of higher experimental binding affinity.

## 6 Discussion

### 6.1 Influence of the embedding

Our results demonstrate that the choice of embedding model has a substantial impact on top-1% retrieval performance. Different retrieval rates upon using ChemBERTa-2 compared to using MolFormer may stem from the different sizes of their pretraining datasets and from different pretraining objectives. MolFormer was trained on 100 million SMILES-encoded molecules, sourced from the complete PubChem database and from a random 10% subset of ZINC.^3^ In comparison, ChemBERTa-2 was pretrained on 77 million molecules exclusively from PubChem. Additionally, MolFormer employs the standard masked language modeling (MLM) objective^39^, whereas ChemBERTa-2 is available in two variants: one trained using MLM and another using a multi-task regression (MTR) objective, which involves predicting RDKit-derived molecular properties from SMILES representations. In this study, we have utilized the MTR version of ChemBERTa-2.

Recently, Sultan *et al*. ^49^ suggested that the MTR objective achieves superior results in property prediction tasks. The binding affinity depends, in addition to properties such as ligand hydrophobicity, also on structural information such as the linker length between two chemical subgroups and packing properties of interface atoms.^56,75^ We hypothesize that subtle differences between the structural properties of molecules are not necessarily reflected in some chemical properties and therefore can lead to similar embeddings of these molecules. As a result, accurate prediction of binding free energies using simple linear regression is nearly impossible. The MLM objective on the other hand is not biased towards embeddings that encode domain specific knowledge (such as physicochemical properties of molecules), but rather implicitly learns the structure of molecules and possibly information relevant for the task at hand. It is important to note that additional protein targets must be tested to draw a definitive conclusion, as the observed results may also originate from MolFormer embeddings more effectively capturing ligand characteristics that may be particularly relevant to the MCL1 protein target and thereby resulting in the observed significantly higher top-1% retrieval rates.

### 6.2 Linear model versus Random Forest

Our findings indicate that a simple Bayesian linear regression model consistently outperforms the Random Forest (RF) model across all datasets and embedding models. Following the approach of Graff *et al*. ^21^ and Cao *et al*. ^25^, the RF model estimates predictive uncertainty as the standard deviation across the outputs of its individual decision trees. As previously reported in both studies, the RF model yields better performance when its mean prediction is used directly as the acquisition function, rather than relying on uncertainty-based strategies such as Expected Improvement (EI) or Upper Confidence Bound (UCB). We therefore attribute the weaker performance of the RF model to its naive uncertainty estimates. Investigating alternative uncertainty estimators^76–78^ represents a promising direction for future research. In contrast, the standard deviation provided by the Bayesian linear regression model captures both the uncertainty due to measurement noise and the uncertainty in the model parameters. ^73^ We assume that this dual consideration acts as an implicit regularizer, helping to mitigate overfitting.

### 6.3 Batch acquisition versus one-at-a-time acquisition

Across all datasets, we observe that selecting a single molecule per iteration consistently results in higher retrieval rates, regardless of the applied embedding or regression model, in particular during the early stages of learning.

Across all datasets, selecting a single molecule per iteration consistently yields higher retrieval rates, especially during early learning, regardless of the embedding or regression model used (Figs. 5 and 6). This is likely due to more frequent model updates, which enhance predictive performance. Unlike batched acquisition, one-at-a-time acquisition allows the model to retrain after each new data point, which is particularly advantageous when training data is still scarce.

While one-at-a-time acquisition improves early retrieval, it limits parallelization because each selection depends on the outcome of the previous simulation. In contrast, batched strategies enable concurrent evaluations, improving resource utilization on parallel architectures and overall throughput. Future work could explore dynamic batching, where new samples are incorporated as soon as computations are complete, to balance frequent updates with parallel efficiency.

### 6.4 Irregular Retrieval Patterns in MMGBSA Compared to Docking

When relying on docking scores, the top-1% retrieval rate improves steadily, plateauing after approximately 2000 iterations (Figure 6) and having reached a high retrieval rate. In contrast, MMGBSA scores often exhibit an early plateau in retrieval rate, sometimes as soon as after 1,000 iterations, before showing a sudden improvement in later stages and eventually stabilizing around 3,000 iterations (Figure 6). It is to note, that such intermediate plateaus tend to occur only for some starting molecules (Figure 8). By choosing different initial datapoints, we can recover similarly steady improvement (Figure 8). This suggests that, instead of relying on a single initial sample, starting with a more diverse sample, potentially trading the increased initial cost for potentially avoiding plateaus in later iterations.

This plateau suggests that, upon learning MMGBSA scores, the model may get trapped in a specific region of the chemical space before eventually identifying a new set of molecular properties that help to escape to other promising regions. Notably, this initial plateau is not observed when using the batched acquisition strategy, presumably because the sampling is too coarse.

Although higher top-1% retrieval rates are achieved when using docking scores, it is important to consider how well these scores actually reflect true binding affinity. To assess this, we have compared predicted binding affinities, based on both docking and MMGBSA scores, with experimentally measured values for a set of known binders (see Figure 2 (a) and Figure 2 (b)). Our analysis reveals that MMGBSA scores exhibit a much stronger correlation with the experimental data (Pearson: 0.610) compared to docking scores (Pearson: 0.177). Furthermore, Figure 2 (c) shows no apparent correlation between MMGBSA and docking scores across our dataset. Notably, the sets of top-1% binders identified by each scoring method differ significantly, with only a 1.6% overlap (represented by the green region in the figure). This may come as a result of MMGBSA accounting for the dynamics of both protein and ligand, in contrast AutoDock Vina relies on simplified empirical scoring functions.^5^ We hypothesize that this makes the relationship between ligand structure and predicted binding affinity more complex, making it more difficult to capture by machine learning models. This added complexity may help explain the observed irregular improvement of top-1% retrieval rates over the iterations.

Moreover, the embedding models used in this study work solely with the SMILES representation of molecules as input and therefore lack explicit information about their three-dimensional structures and spatial orientation within the protein’s binding pocket. Such structural context may be critical for accurately capturing the interactions that influence MMGBSA scores. The absence of this information could be a key factor limiting the model’s ability to achieve top-1% retrieval rates with MMGBSA scores comparable to those observed with docking scores.

## 7 Conclusion and Future Work

We presented an active learning approach that combines pretrained embedding models with lightweight Bayesian regression models to predict binding affinities. In contrast to previous methods that require fine-tuning of the entire model for each new batch of data, our strategy greatly reduces computational overhead by updating only the parameters of the regression head. This design enables the use of smaller query batches, facilitating a more responsive feedback loop and ultimately leading to improved retrieval of top binders.

Given the high computational demands of MD-based binding free energy estimations, existing datasets typically contain only a few thousand data points. In this study, we computed binding free energies with MMGBSA for approximately 60,000 ligands targeting MCL1. To test our active learning framework, we compared its performance upon using MMGBSA scores relative to using docking scores. After querying only 6% of the dataset, the model successfully identified 79.9% of the top-1% ligands according to MMGBSA, laying the ground for large-scale virtual screenings with MMGBSA-level accuracy. Additionally, our results indicate that docking scores are easier to predict than MMGBSA scores, which may be attributed to the more dynamic transformation process of the protein and ligand when conducting the MD simulation for MMGBSA. We hypothesize that enriching SMILES-based embeddings with additional information, particularly regarding ligand pose within the binding pocket, may help address this information gap and alleviate the observed performance difference. While MMGBSA scores are more challenging for the model to learn, they exhibit a stronger correlation with experimental binding affinities than docking scores in the subset of experimentally known MCL1 binders. This suggests that top binders selected based on MMGBSA may represent more promising candidates than those selected with docking. Based on our results, this active learning pipeline is now ready to incorporate additional scores into the learning loop such as affinities from free energy perturbation simulations or from wet lab experiments.

## 8 Conflicts of interest

There are no conflicts to declare.

## Acknowledgements

We thank Afnan Sultan for helpful discussions. This research was funded in part by the European Union’s Horizon 2020 Research and Innovation Program under Marie Skłodowska Curie Grant 860592 and in part by the Deutsche Forschungsgemeinschaft (DFG, German Research Foundation; grant INST 256/539-1).

## Data and Software Availability

All relevant data and software is provided via our GitHub repository https://github.com/uds-lsv/bayesian-optimization-mmgbsa. The dataset and the ligand poses are also available at https://doi.org/10.5281/zenodo.17579183.

## Supporting Information

**Figure S1.**
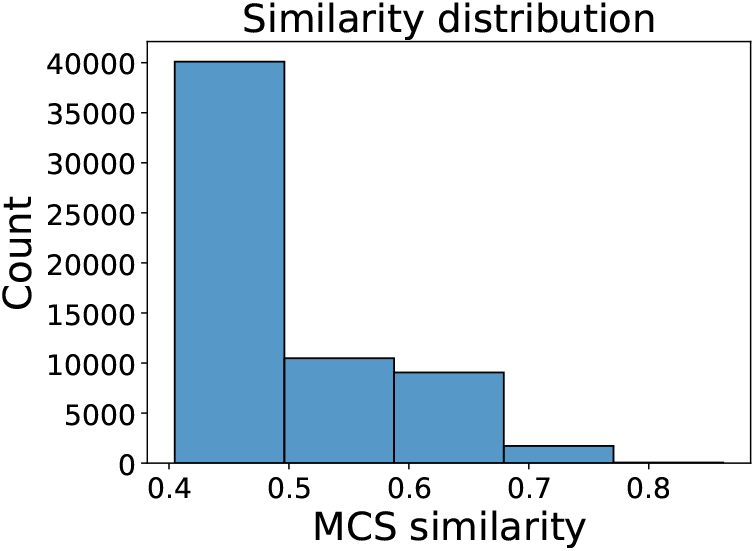
Histogram of MCS-Tanimoto similarity to the query compound. The dataset was screened with a minimum similarity threshold of 0.4. Most compounds show low similarity values, typical for chemical datasets.

**Figure S2.**
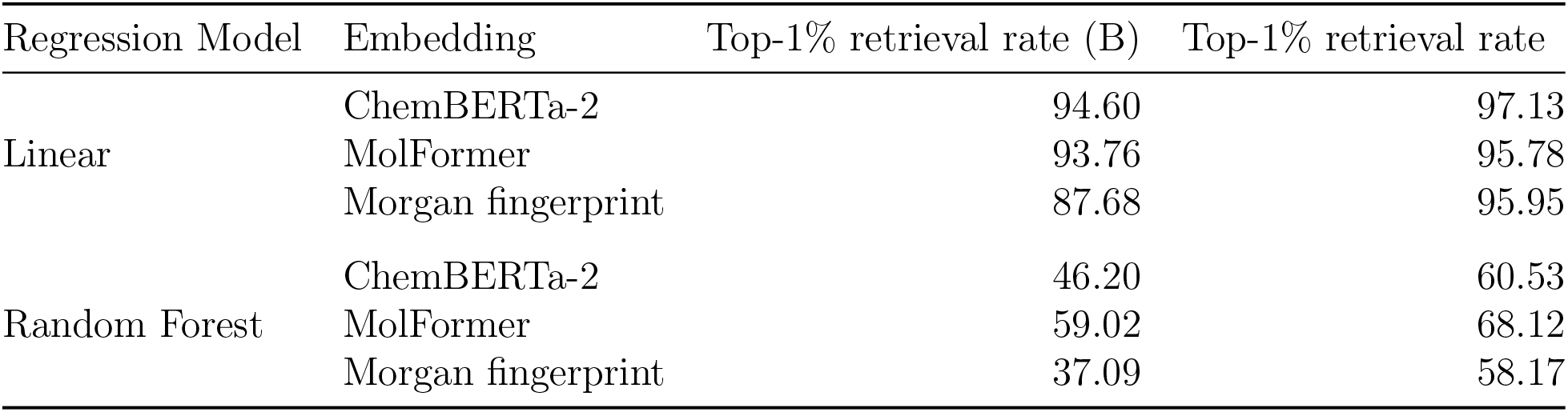
Retrieval rates of all embedding-regression model combinations on the `MCL1-Docking` data. (B) indicates batched querying of 1% of the dataset for 6 iterations.

**Figure S3.**
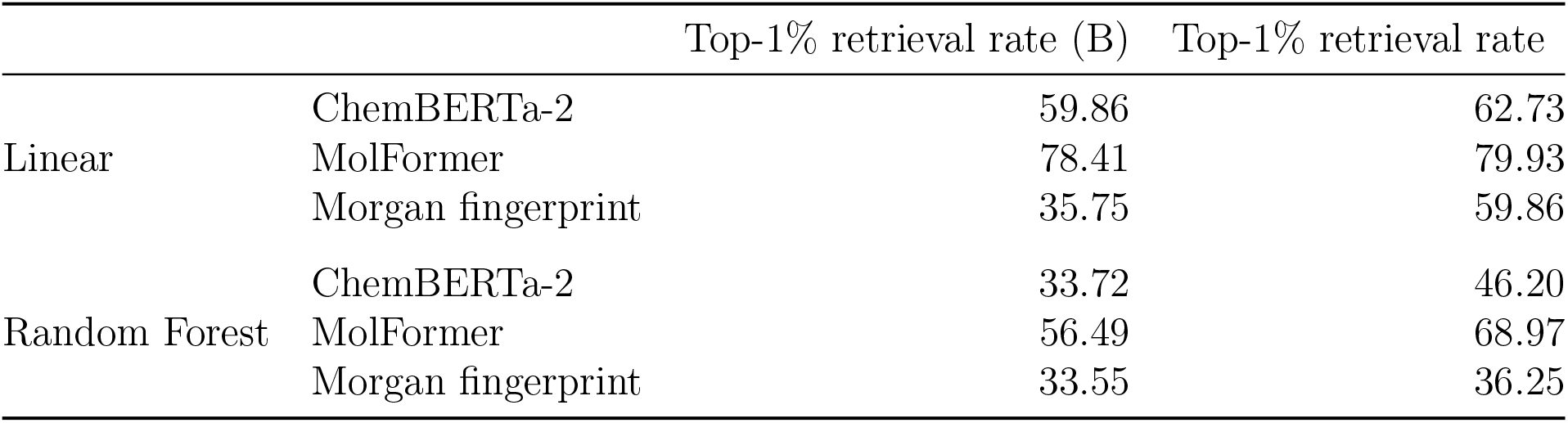
Retrieval rates of all embedding-regression model combinations on the `MCL1-MMGBSA` data. (B) indicates batched querying of 1% of the dataset using 6 iterations.

**Figure S4.**
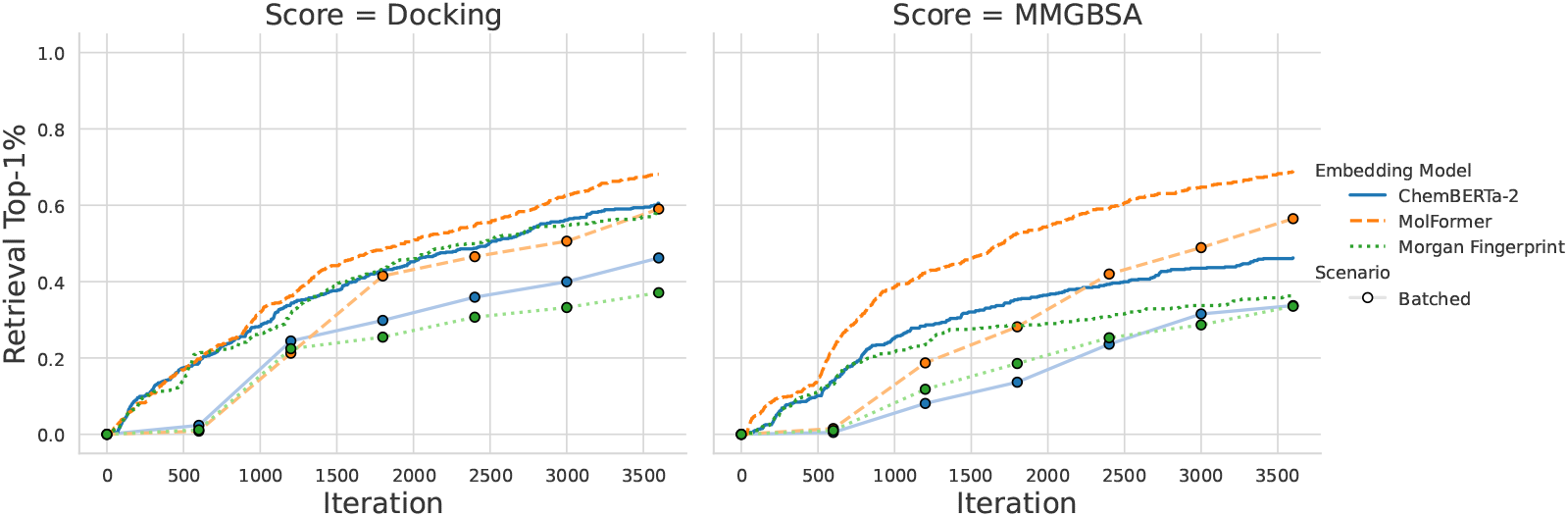
Retrieval of top-1% molecules on docking and MMGBSA scores using a Random Forrest instead of a linear regression model as presented in Fig. 6.

**Figure S5.**
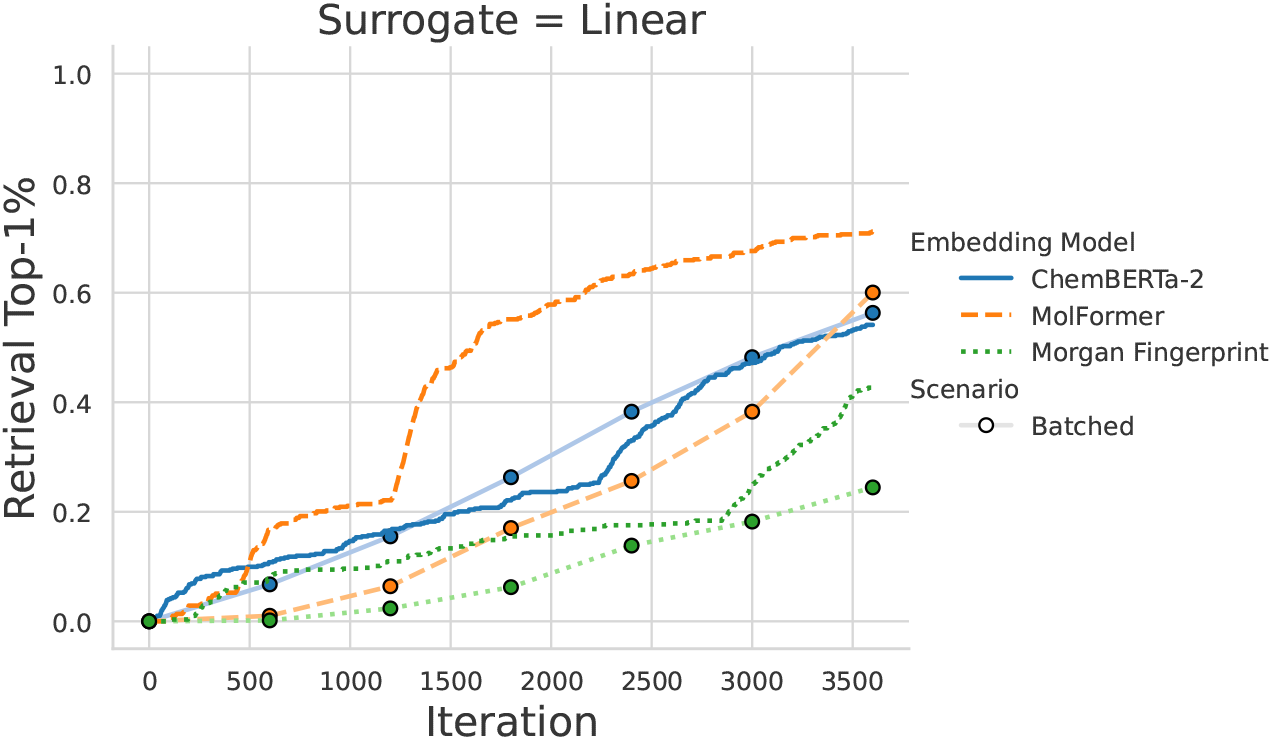
Retrieval of top-1% molecules on MMPBSA scores using a linear regression model.

**Figure S6.**
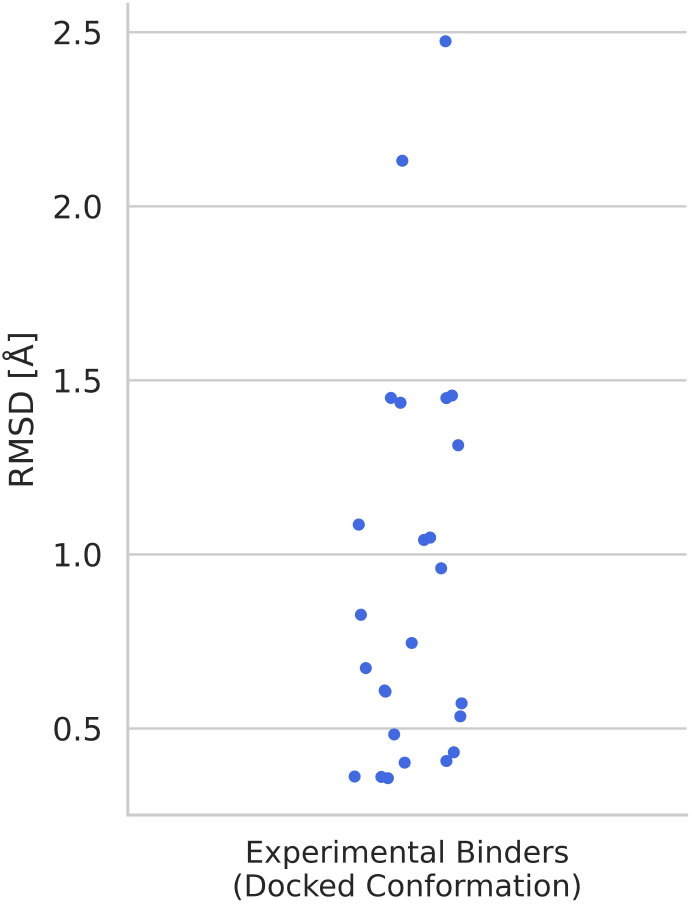
Distribution of root mean-square deviation (RMSD) values for the maximum common substructure (MCS) between docked poses of experimental binders^56^ and the reference ligand in the MCL1 crystal structure^56^. Each point represents the RMSD of an individual ligand, calculated using only heavy atoms. All ligands were docked into the same MCL1 protein structure, and RMSD values were computed without explicit alignment to the reference ligand. The generally low RMSD values indicate that AutoDock Vina consistently positioned the experimental binders in conformations closely resembling that of the crystallographic reference ligand.

**Figure S8.**
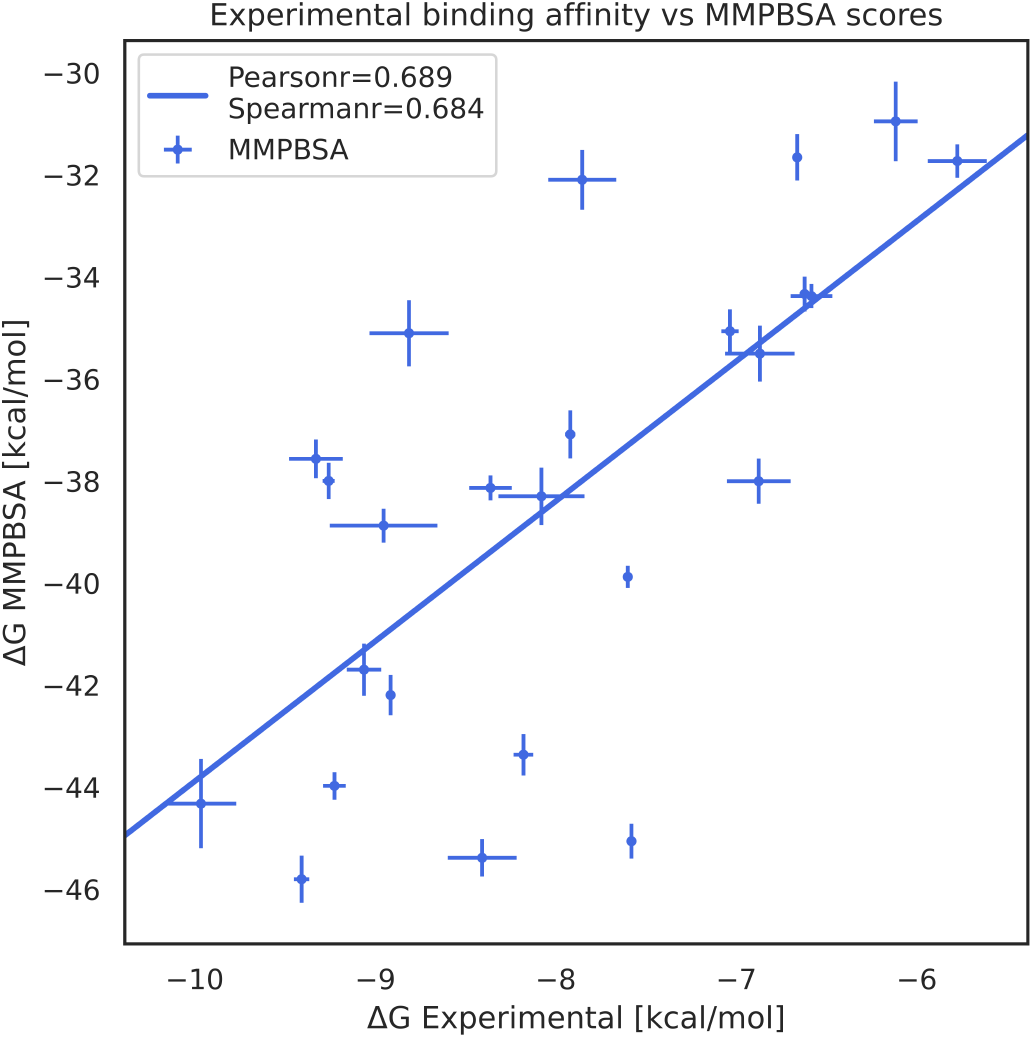
Hexbin plots of the normalized activity difference binned by distance in MolFormer embedding space. For the same distance, the difference in the activity value for MMGBSA is generally larger than for Vina, indicating a less smooth energy surface.

**Figure S7.**
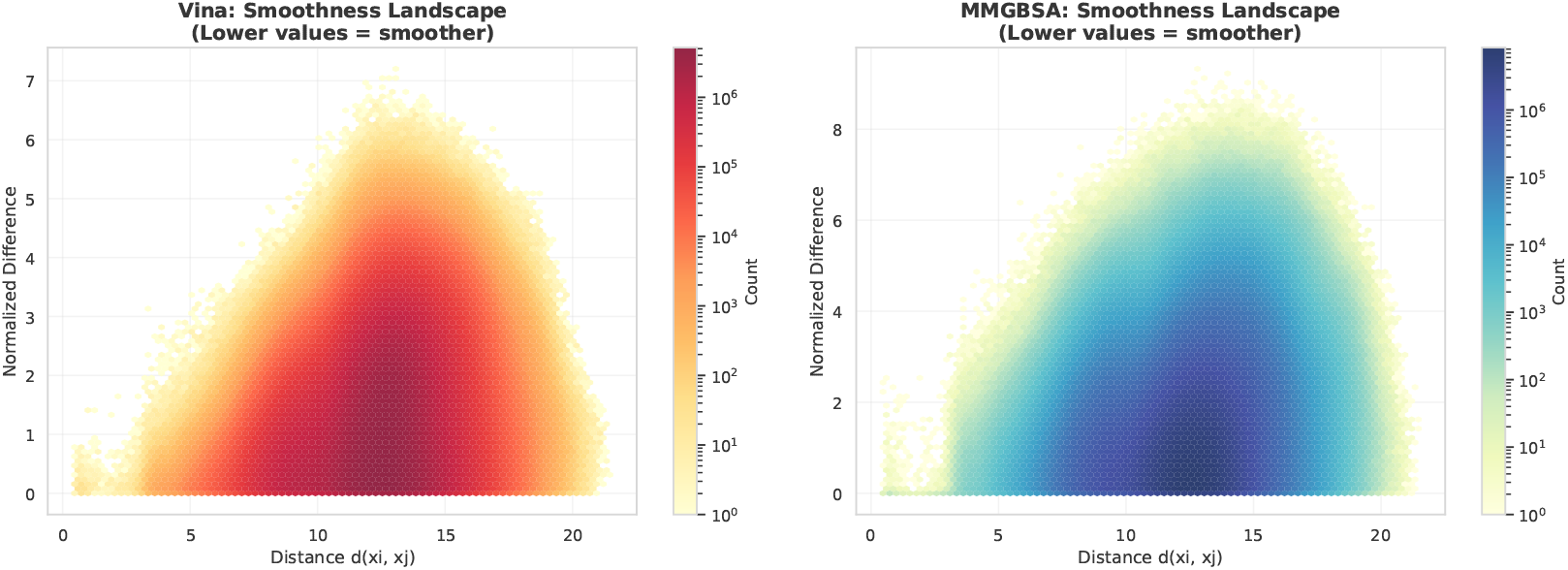
Correlation between MMPBSA (instead of MMGBSA) scores with experimental binding affinities for experimental binders. Pearson and Spearman correlation coefficients are slightly higher compared to results based on MMGBSA (compare with Fig. 2).

## Notes

### Competing Interest Statement

The authors have declared no competing interest.

### Summary of Updates

Minor clarifications: * description of the dataset of 60000 molecules simulated * discussion of the MMGBSA affinity landscape * various other minor clarifications

